# First high-quality reference genome of *Amphicarpaea edgeworthii*

**DOI:** 10.1101/2020.09.22.306811

**Authors:** Tingting Song, Mengyan Zhou, Yuying Yuan, Jinqiu Yu, Hua Cai, Jiawei Li, Yajun Chen, Yan Bai, Gang Zhou, Guowen Cui

## Abstract

*Amphicarpaea edgeworthii*, an annual twining herb, is a widely distributed species and an ideal model for studying complex flowering types and evolutionary mechanisms of species. Herein, we generated a high-quality assembly of *A. edgeworthii* by using a combination of PacBio, 10× Genomics libraries, and Hi-C mapping technologies. The final 11 chromosome-level scaffolds covered 90.61% of the estimated genome (343.78 Mb), which is the first chromosome-scale assembled genome of an amphicarpic plant. These data will be beneficial for the discovery of genes that control major agronomic traits, spur genetic improvement of and functional genetic studies in legumes, and supply comparative genetic resources for other amphicarpic plants.

## Introduction

In nature, the distribution of key resources required for plant growth is often uneven. Plants growing in unstable habitats, with limited supplies of mineral nutrients, water, or light, frequent soil interferences, and large environmental fluctuations, undergo adaptive evolution to improve their survival(Jackson and Caldwell, 1993; Pearcy and Caldwell, 1994). Some plant species that bear 2 or more heteromorphic flowers also bear heteromorphic fruits (seeds). Amphicarpy is a phenomenon in which a plant produces both aerial and subterranean flowers and simultaneously bears both aerial and subterranean fruits (seeds) on aerial- and subterranean-stems, respectively (Cheplick, 1987; Koontz et al., 2017; Schnee and Waller, 1986). This phenomenon is observed in at least 67 herbaceous species (31 in Fabaceae) in 39 genera and 13 families of angiosperms, as reported by Zhang et al(Zhang et al., 2020a). Amphicarpy is an important part of plant adaptive evolution, in which angiosperms generally display a special type of fruiting pattern and different fruit (seed) types also exhibit various dormancy and morphological features. This type of fruiting mode is crucial for the ecological adaptation of plants that have evolved through natural selection because it reduces competition among siblings within the population, maintains and increases the population size in situ, and increases the adaptability and evolutionary plasticity of the species(Hidalgo et al., 2016; Sadeh et al., 2009).

*Amphicarpaea edgeworthii*, an annual twining herb, belongs to the Fabaceae family, which is a large and economically valuable family of flowering plants(Zhang et al., 2017; Zhang et al., 2006). In this plant species, 3 types of flowers (fruits) can grow in the same plant, and aerial chasmogamous flowers are produced only during summer(Zhang et al., 2006; Zhang et al., 2005). This species may offer an attractive model for examining gene regulatory networks that control chasmogamous and cleistogamous flowering in plants. However, the mechanism of flower development in amphicarpic plants, particularly in legumes, is sparsely known. The present study was an attempt to enhance our understanding on the reproductive biology and the ultimate evolutionary mechanism in amphicarpic plants. We performed whole-genome sequencing of *A. edgeworthii*, which is the first plant in the legume family and also the first amphicarpic species subjected to sequencing. This reference genome represents a precious foundation for further understanding on agronomics and molecular breeding in *A. edgeworthii*.

## Result

*A. edgeworthii* has a diploid genome (2n = 2x = 22) (Supplementary Figure 1). We estimated the genome size of *A. edgeworthii* to be 360.91 Mb using 17-mer(Supplementary Table 1 and Figure 2). We sequenced the genome of *A. edgeworthii* by using a combination of PacBio, Illumina, and 10× Genomics libraries that resulted in the generation of a 343.78-Mbp genome (contig N50 length = 1.44 Mb, scaffold N50 length = 2.4 Mb; Table 1 and Supplementary Table 2, 3 and 4). Finally, we assembled a chromosome-level genome by using Hi-C technology. We used a total of 5.27 million reads from Hi-C libraries and mapped approximately 90.61% of the assembled sequences to 11 pseudochromosomes, with the longest scaffold length of 32.05 Mb (Table 1 and Figure 1). Results indicate that the *A. edgeworthii* genome was adequately covered by the assembly.

**Table 1.**
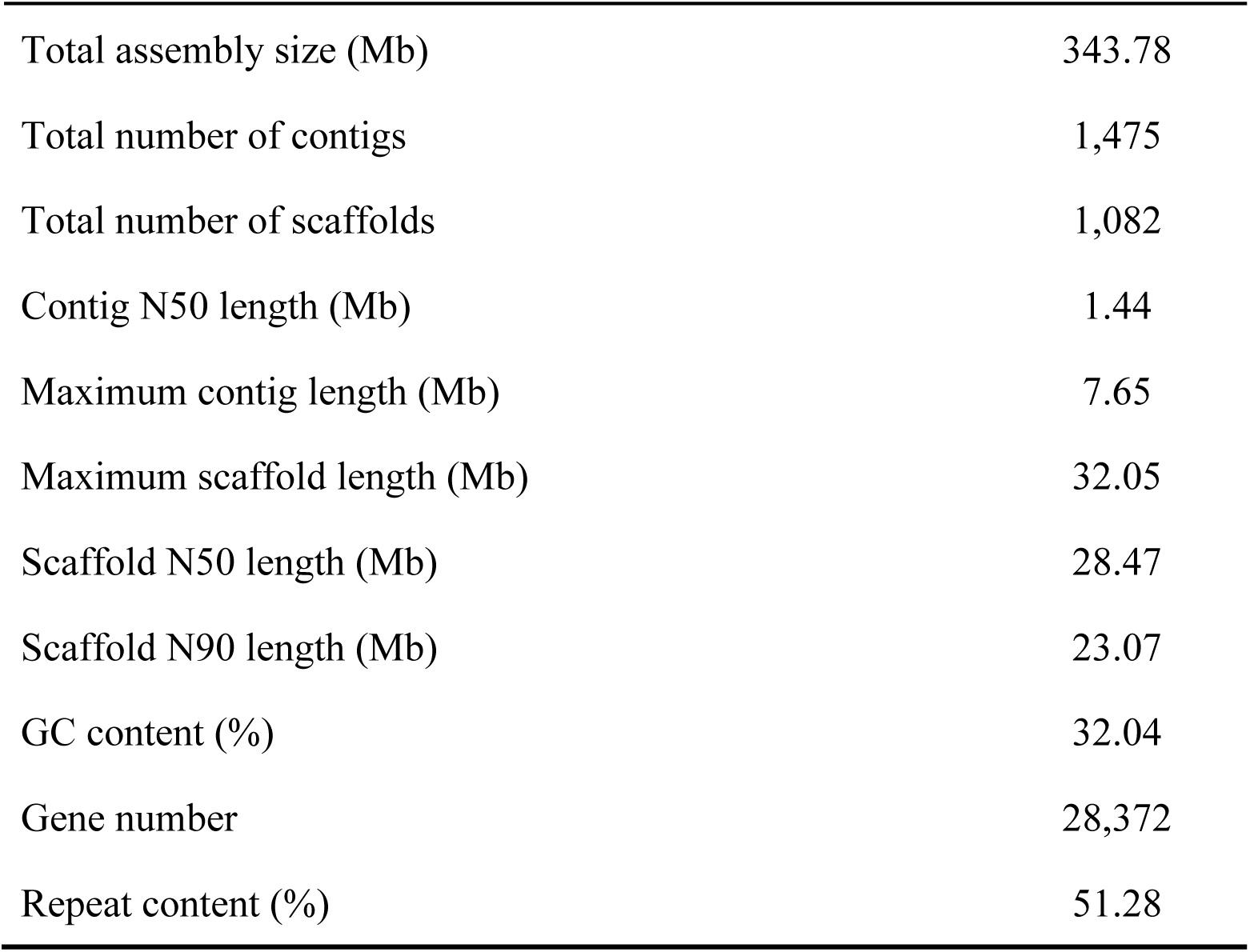
Statistics of the *A. edgeworthii* genome assembly.

|  |  |
| --- | --- |
| Total assembly size (Mb) | 343.78 |
| Total number of contigs | 1,475 |
| Total number of scaffolds | 1,082 |
| Contig N50 length (Mb) | 1.44 |
| Maximum contig length (Mb) | 7.65 |
| Maximum scaffold length (Mb) | 32.05 |
| Scaffold N50 length (Mb) | 28.47 |
| Scaffold N90 length (Mb) | 23.07 |
| GC content (%) | 32.04 |
| Gene number | 28,372 |
| Repeat content (%) | 51.28 |

**Figure 1.**
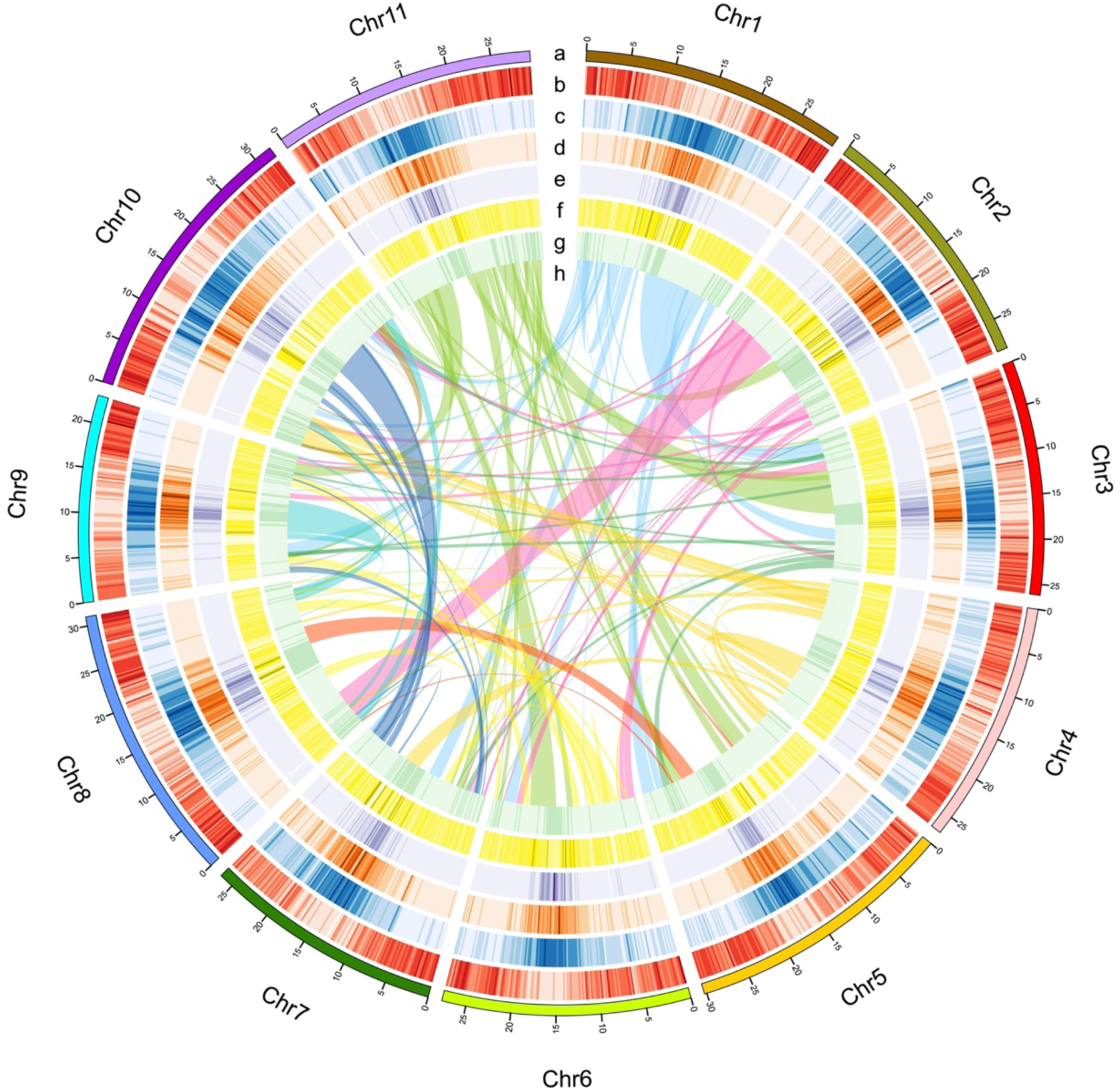
Characteristics of the *A. edgeworthii* genome: **a**. Chromosome length; **b**. Gene density per chromosome; **c**. Repeat density; **d**. LTR_Copia density; **e**. LTR_Gypsy density; **f**. SNP density in 5 populations; **g**. Distribution of GC content; **h**. Intra-genome collinear blocks connected. All statistics are computed for windows of 200 kb.

We evaluated the completeness of the genome assembly by mapping the Illumina paired-end reads to our assembly through Burrows–Wheeler Alignor (BWA) (Li and Durbin, 2009) with 98.70 % of mapping rate and 94.04 % of coverage (Supplementary Table 5). Then, we used both Core Eukaryotic Gene Mapping Approach (CEGMA) (Parra et al., 2007) and Benchmarking Universal Single-Copy Orthologs (BUSCO) (Simão et al., 2015) to access the integrity of the assembly. In the CEGMA assessment, 238 (95.97%) of 248 core eukaryotic genes were assembled (Supplementary Table 6). Furthermore, 93.4% complete single-copy BUSCOs were detected, which indicates that the assembly was complete (Supplementary Table 7). Overall, the results indicate that a high-quality assembly was generated.

Repeat sequences comprise 51.28% of the assembled genome, with transposable elements (TEs) being the major component (Supplementary Table 8). Among the TEs, long terminal repeats (LTRs) were the major component (29.32%) (Supplementary Table 9). We combined de novo prediction, homology search, and mRNA-seq assisted prediction to predict genes in the *A. edgeworthii* genome, and we obtained 28,372 protein-coding genes (97.2% were annotated) (Supplementary Tables 10 and 11). Additionally, we identified 2,260 non-coding RNAs, including 471 miRNAs, 701 transfer RNAs, 266 ribosomal RNAs, and 822 small nuclear RNAs (Supplementary Table 12).

## Discussion

In this study, we constructed a high-quality chromosome-level genome assembly for *A. edgeworthii* by combining the long-read sequences from PacBio with highly accurate short reads from Illumina sequencing and using Hi-C technology for super-scaffolding. The assembly of *A. edgeworthii* adds to the growing genomic information on the Fabaceae family. As the first species in the Amphicarpaea genera of the Fabaceae family to be sequenced, *A. edgeworthii* exhibits a range of specific biological features, such as 3 types of flowers and 3 types of fruit (seed) simultaneously in the same plant. It has maintained an essential evolutionary position in the tree of life(Goodwillie et al., 2005; Silvertown et al., 2001). According to our analysis, these genomic data will serve as valuable resources for future genomic studies on amphicarpic plants and molecular breeding of soybean.

To date, no genomic data for the amphicarpic plants are available. Therefore, the sequencing of the entire *A. edgeworthii* genome will facilitate future investigations on the phylogenetic relationships of the flowering (seed) plants. The accessibility of the *A. edgeworthii* genome sequence makes feasible the investigation of deep phylogenetic questions on angiosperms and the determination of genome evolution signatures and the genetic basis of interesting traits.

## Method

### Plant material and genome survey

For genome sequencing, we collected fresh young leaves of *A. edgeworthii* species distributed in Heilongjiang Province (45.80°N, 126.53°E), China. The karyotype analysis of the plant species revealed a karyotype of 2n = 2x = 22, with uniform and small chromosomes(Wolny et al., 2013) (Supplementary Figure 1).

We extracted DNA from the fresh leaves of *A. edgeworthii* by using a DNAsecure Plant Kit(TIANGEN, Biotech, China) and then purified and concentrated, high-quality DNA was broken into random fragments, and Illumina paired-end library with 350-bp size was constructed and was sequenced using a Illumina HiSeq X-ten platform.

To estimate the *A. edgeworthii* genome size, high-quality short-insert reads (350-bp size) were used to extract the 17-k-mer sequences by using sliding windows. The frequency of each 17-k-mer was calculated and is presented in Supplementary Figure 2. Genome size was calculated by using the following formula:

Genome size = total k-mer numbers/k-mer depth

The revised genome size was calculated after excluding the erroneous k-mers (Supplementary Tables 1).

### Library construction, genome sequencing, assembly, and evaluation

To construct long-insert libraries, the SMRTbell libraries were constructed by following the standard protocol as per the manufacturer’s instructions (PacBio Biosciences). Genomic DNA was broken into fragments of 15 kb–40 kb in size and the large fragments were enriched and enzymatically repaired and converted into SMRTbell libraries. The SMRTbell libraries were sequenced using a PacBio Sequel platform.

The linked read sequencing libraries of 10× Genomics GemCode platform(Weisenfeld et al., 2017) were sequenced with 350-bp size by using an Illumina HiSeq X-ten platform.

Fresh leaves were plucked from the plant and chromatin in the samples were cross-linked to DNA and fixed. A chromatin interaction mapping (Hi-C) library with 350-bp size was constructed for sequencing using Illumina HiSeq X-ten.

We used FALCON software(Chin et al., 2016) for de novo assembly of PacBio SMRT reads (Supplementary Tables 2 and 3). The longest coverage of subreads were selected as seeds for assembly after pairwise comparisons of all reads for error correction with default parameters. The error-corrected SMRT reads were aligned to each other to construct string graphs. After initial assembly, the produced contigs were polished using Quiver(Chin et al., 2013) with default parameters. The first round of error correction was performed using Illumina paired-end reads by Pilon(Walker et al., 2014). Subsequently, the scaffolding was performed using 10× Gscaff v2.1 with 10× genomics data, and the genome was upgraded by PBjelly(English et al., 2012). The second round of error correction was performed using Illumina paired-end reads by Pilon(Walker et al., 2014). The Hi-C data were mapped to the original scaffold genome by using BWA v0.7.7(Li and Durbin, 2009), and only the reads with unique alignment positions were extracted to construct a chromosome-scale assembly by using the Ligating Adjacent Chromatin Enables Scaffolding In Situ(LACHESIS) tool(Burton et al., 2013) (Supplementary Table 3).

### Genome annotation

We used RepeatModeler, RepeatScout(Tarailo and Chen, 2009), Piler(Edgar and Myers, 2005), and LTR_FINDER(Xu and Wang, 2007) to develop a de novo transposable element library. RepeatMasker(Tarailo and Chen, 2009) was used for DNA-level identification in the Repbase and de novo transposable element libraries. Tandem repeats were ascertained in the genome by using Tandem Repeats Finder(Benson, 1999). RepeatProteinMask(Tarailo and Chen, 2009) was used to conduct WU-BLASTX searches against the transposable element protein database. Overlapping TEs belonging to the same type of repeats were integrated (Supplementary Tables 8 and 9).

To predict protein-coding genes in the *A. edgeworthii* genome, we used homolog-based prediction (using *A. duranensis, C. arietinum, G. max, M. truncatula, P. vulgaris*, and *T. pratense* gene sets), de novo prediction (using Augustus v.3.0.2(Stanke and Morgenstern, 2005), Genescan v.1.0(Aggarwal and Ramaswamy, 2002), GeneID(Parra et al., 2000), GlimmerHMM v.3.0.2(Majoros et al., 2004), and SNAP(Korf, 2004) programs) and transcriptome-based prediction (using 5 tissue RNA sequencing data). A weighted and non-redundant gene set was generated using EVidenceModeler (EVM) (Brian et al., 2008), which merged all the genes models that were predicted using the aforementioned approaches. Along with the transcript assembly, the Program to Assemble Spliced Elements(Haas et al., 2003) was used to adjust the gene models generated using EVM (Supplementary Table 10).

Functional annotation of protein-coding genes was evaluated using BLASTP (E-value ≤ 1E-05) against 2 integrated protein sequence databases, SwissProt(Bairoch and Apweiler, 2000) and NCBI non-redundant protein database. Protein domains were annotated by searching InterPro v.32.0, which included Pfam, PRINTS, PROSITE, ProDom, and SMART databases, by using InterProScan v.4.8(Mulder and Apweiler, 2007). GO(Ashburner et al., 2000) terms for each gene were obtained from the corresponding InterPro descriptions. The pathways in which the gene might be involved were assigned using BLAST searches against the KEGG database(Kanehisa and Goto, 2000), with an E-value cutoff of 1E-05 (Supplementary Table 11).

The tRNA genes were predicted using tRNAscan-SE software(Lowe and Eddy, 1997). The miRNA and snRNA fragments were identified using INFERNAL software(Nawrocki et al., 2009) against the Rfam(Griffiths et al., 2005) database. The rRNA were identified using BLASTN searches (E-value ≤ 1E-10) against the plant rRNA database (Supplementary Table 12).

## Acknowledgement

This research work was funded by the National Key R&D Project of China (2016YFC0500607). We are also grateful to Ms. Mei Zhang, Mr. Bo Zhao and Ms. Junyan Li (Novogene Bioinformatics Institute, Beijing China) for providing helps in the project.

## Author contributions

G.W.C.and T.T.S. conceived and designed the project and the strategy; T.T.S, Y.Y.Y, J.Q.Y and J.W.L. collected and cultured the plant material, DNA/RNA preparation, library construction and sequencing; T.T.S and M.Y.Z worked on genome assembly and annotation, comparative and population genomic analyses, and transcriptome sequencing and analysis; G.W.C., T.T.S, M.Y.Z, H.C., Y.J.C, Y.B, and G.Z contributed substantially to revisions. All authors commented on the manuscript.

