## Supplemental Information for "First high-quality reference genome of *Amphicarpaea edgeworthii*"

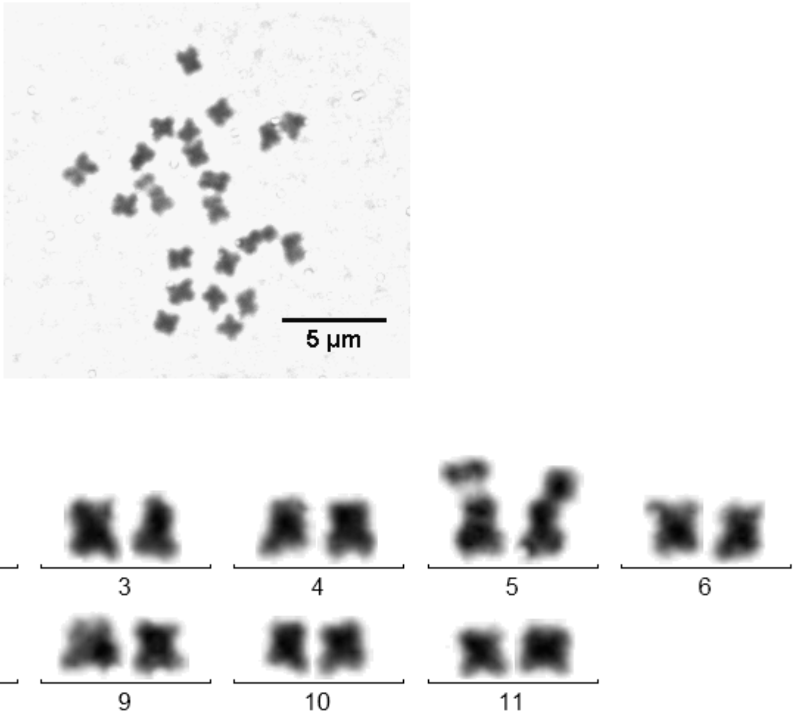


**Supplementary Figure 1. Mitotic metaphase chromosomes of the sequenced individual of *A.edgeworthii*.** The chromosome number is 22, indicating the sequenced individual is diploid. **Related to Figure 1.**

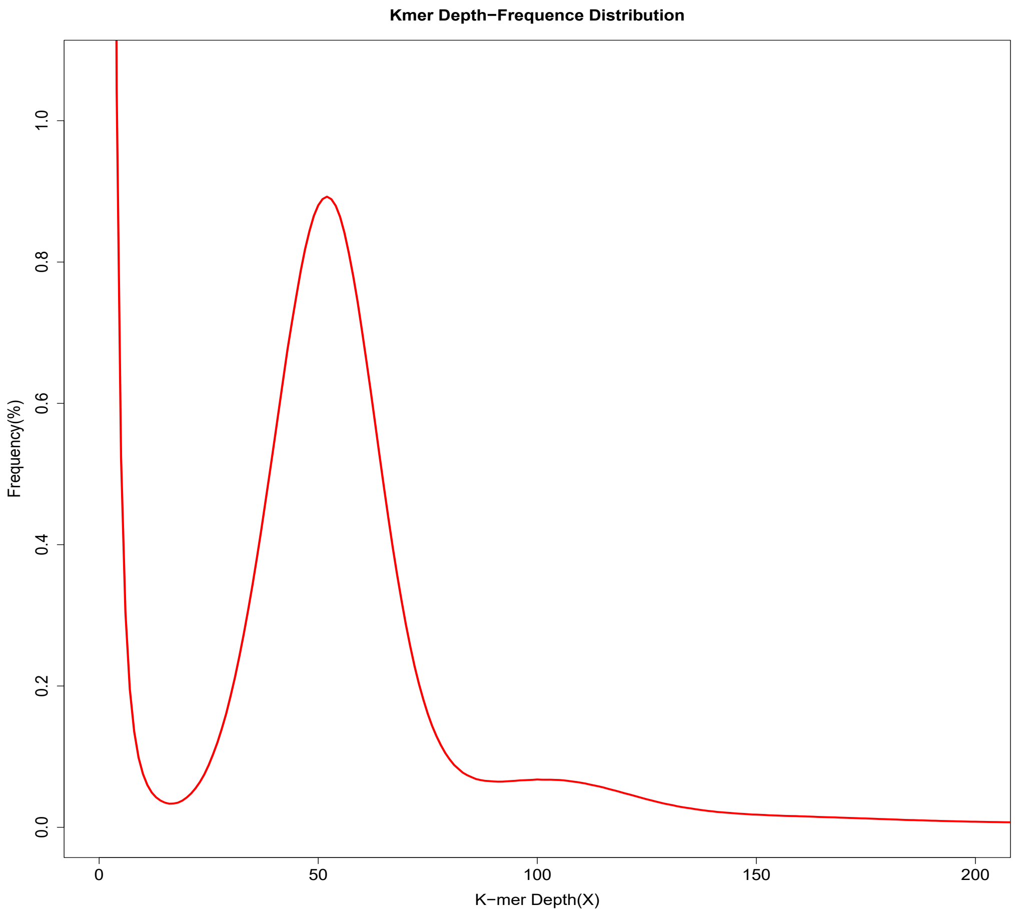


**Supplementary Figure 2. 17-Kmer depth distribution of the *A.edgeworthii* genome sequencing reads.** The depth of peak was at 53. **Related to Figure 1.**

**Supplementary Table 1. Statistics for K-mer analysis. Related to Figure 1.**

| K-mer | K-mer Number | K-mer Depth | Genome Size (Mb) | Heterozygous Ratio (%) | Repeat(%) |
|---|---|---|---|---|---|
| 17 | 19,453,880,408 | 53 | 360.91 | 0.30 | 48.14 |

**Supplementary Table 2. Sequencing data generated from different platforms or strategies. Related to Figure 1.**

| Platform | Insert size | Read length (bp) | Raw data (G) | Sequence coverage (X) # |
|---|---|---|---|---|
| Illumina | 350bp | 150 | 103.96 | 288.05 |
| PacBio | -- | -- | 37.32 | 103.41 |
| 10X Genomics | -- | 150 | 138.63 | 384.11 |
| Hi-C | 350 bp | 150 | 45.38 | 125.76 |
| Total | -- | -- | 325.29 | 901.31 |

### The estimated genome size is ~360.91 Mb, and sequence coverage = Total raw bases/360.91 Mb.

**Supplementary Table 3. Statistics of the assembly of the *A.edgeworthii* genome. Related to Figure 1.**

| Sample ID | Length | | Number | |
|---|---|---|---|---|
| | Contig (bp) | Scaffold (>100 bp) | Contig | Scaffold (>100 bp) |
| Total | 343,740,538 | 343,779,838 | 1,475 | 1,082 |
| Max | 7,653,304 | 32,051,318 | - | - |
| Number>=100 | - | - | 1,475 | 1,082 |
| Number>=2000 | - | - | 1,435 | 1,042 |
| N50 | 1,444,032 | 28,474,477 | 67 | 6 |
| N60 | 1,113,772 | 27,686,701 | 93 | 7 |
| N70 | 831,149 | 26,642,412 | 129 | 9 |
| N80 | 448,254 | 26,200,017 | 183 | 10 |
| N90 | 111,078 | 23,069,742 | 315 | 11 |

Note: N50 refers to the length of sequence equal to or greater than the half of total sequences length.

**Supplementary Table 4. Statistics for GC content. Related to Figure 1.**

| | Number | % of genome |
|---|---|---|
| **A** | 116,788,363 | 33.97% |
| **T** | 116,832,048 | 33.98% |
| **C** | 55,078,855 | 16.02% |
| **G** | 55,041,272 | 16.01% |
| **N** | 39,300 | 0.01% |
| **Total** | 343,779,838 | -- |
| **GC** | 110,120,127 | 32.04% |

Note: GC content of the genome without N

**Supplementary Table 5. Statistics of paired-end reads mapping. Related to Figure 1.**

| | Sample ID | Percentage |
|---|---|---|
| Reads | Mapping rate (%) | 98.70 |
| | Average sequencing depth | 217.51 |
| | Coverage (%) | 94.04 |
| Genome | Coverage at least 4X (%) | 93.06 |
| | Coverage at least 10X (%) | 92.21 |
| | Coverage at least 20X (%) | 91.38 |

**Supplementary Table 6. CEGMA results of *A.edgeworthii* genome. Related to Figure 1.**

| Species | Complete | | Complete + partial | |
|---|---|---|---|---|
| | Prots | % completeness | Prots | % completeness |
| *A.edgeworthii* | 235 | 94.76 | 238 | 95.97 |

Note: A protein is classified as complete if the alignment of the predicted protein to the HMM profile represents at least 70% of the original KOG domain, otherwise is classified as partial.

**Supplementary Table 7. BUSCO (Benchmarking Universal Single-Copy Orthologs) results of *A.edgeworthii* genome. Related to Figure 1.**

| Species | BUSCO notation assessment results |
|---|---|
| *Amphicarpaea edgeworthii* | C: 93.4% [S: 82.9%,D: 10.5%], F: 1.2%, M: 5.4%, n:1440 |

C: Complete BUSCOs

S: Complete and single-copy BUSCOs

D: Complete and duplicated BUSCOs

F: Fragmented BUSCOs

M: Missing BUSCOs

n: Total BUSCO groups searched

**Supplementary Table 8. The prediction of repeats elements in *A. edgeworthii* genome. Related to Figure 1.**

| Type | Repeat Size (bp) | Percent (%) |
|---|---|---|
| TRF | 12,887,689 | 3.73 |
| Repeat Masker | 136,769,884 | 39.64 |
| Repeat Protein Mask | 37,476,456 | 10.86 |
| Total | 176,927,760 | 51.28 |

**Supplementary Table 9. Categories of TEs predicted in *A. edgeworthii* genome. Related to Figure 1.**

| | | **Repbase + De novo** | | **TE Proteins** | | **Combined TEs** | |
|---|---|---|---|---|---|---|---|
| | | **Length (bp)** | **% in Genome** | **Length (bp)** | **% in Genome** | **Length (bp)** | **% in Genome** |
| DNA transposon | DNA | 20,929,453 | 6.065557 | 4,642,817 | 1.345533 | 25,572,270 | 7.41109 |
| Retrotransposon | LINE | 17,702,640 | 5.130395 | 8,461,973 | 2.452361 | 26,164,613 | 7.582756 |
| | SINE | 103,777 | 0.030076 | 0 | 0 | 103,777 | 0.030076 |
| | LTR | 76,745,004 | 22.241439 | 24,415,265 | 7.075778 | 101,160,269 | 29.317218 |
| Other | Other | 956 | 0.000277 | 0 | 0 | 956 | 0.000277 |
| | Unknown | 31,698,364 | 9.18649 | 0 | 0 | 31,698,364 | 9.18649 |
| | Total | 136,769,884 | 39.637226 | 37,476,456 | 10.861037 | 174,246,340 | 50.498263 |

Note: "Other" refer to the repeats that can be classified by Repeat Masker, but not included by the classes above; "Unknown" refer to the repeats that can’t be classified by Repeat Masker.

**Supplementary Table 10. General statistics of predicted protein-coding genes. Related to Figure 1.**

| | Gene set | Number | Average transcript length (bp) | Average CDS length (bp) | Average exon length (bp) | Average intron length (bp) | Average exons per gene |
|---|---|---|---|---|---|---|---|
| *De novo*# | Augustus | 31,232 | 2,707.40 | 1,105.86 | 229.60 | 419.63 | 4.82 |
| | GlimmerHMM | 48,340 | 5,247.00 | 724.84 | 222.79 | 2,006.82 | 3.25 |
| | SNAP | 52,948 | 2,402.47 | 752.34 | 173.45 | 494.40 | 4.34 |
| | Genscan | 37,398 | 4,536.99 | 901.00 | 198.05 | 1,024.43 | 4.55 |
| | Geneid | 26,795 | 7,340.67 | 1,205.21 | 210.62 | 1,299.24 | 5.72 |
| Homolog$ | *Arachis duranensis* | 24,738 | 2,963.43 | 1,266.57 | 261.28 | 441.03 | 4.85 |
| | *Cicer arietinum* | 26,099 | 2,903.95 | 1,219.39 | 264.54 | 466.71 | 4.61 |
| | *Phaseolus vulgaris* | 34,065 | 2,361.38 | 1,039.08 | 247.95 | 414.92 | 4.19 |
| | *Trifolium pratense* | 25,178 | 2,709.65 | 1,172.09 | 260.70 | 439.81 | 4.50 |
| | *Medicago truncatula* | 35,368 | 2,220.26 | 976.45 | 250.90 | 430.11 | 3.89 |
| | *Glycine max* | 35,088 | 2,478.73 | 1,037.27 | 248.55 | 454.25 | 4.17 |
| RNA_Seq | Cufflinks | 45,660 | 5,335.79 | 2,257.92 | 327.00 | 521.24 | 6.90 |
| | PASA | 86,731 | 2,738.81 | 1,022.99 | 202.66 | 423.90 | 5.05 |
| EVM | | 35,144 | 2,699.21 | 1,042.15 | 224.52 | 455.03 | 4.64 |
| PASA update | | 34,789 | 2,708.97 | 1,065.56 | 227.66 | 446.52 | 4.68 |
| Final set + | | 28,372 | 2,968.57 | 1,152.33 | 224.42 | 439.26 | 5.13 |

### Statistics calculated from the gene set predicted form each method.

$ Statistics calculated from the gene set predicted by homolog proteins from each species.

+ Statistics calculated from the *Amphicarpaea edgeworthii* genome.

**Supplementary Table 11. General statistics of mapping rate to functional database of protein-coding genes. Related to Figure 1.**

| **Database** | | **Annotated Number** | **Annotated Percent (%)** |
|---|---|---|---|
| NR | | 27,565 | 97.2 |
| Swiss-Prot | | 21,711 | 76.5 |
| KEGG | | 20,507 | 72.3 |
| InterPro | All | 23,127 | 81.5 |
| | Pfam | 21,558 | 76.0 |
| | GO | 15,609 | 55.0 |
| Annotated | | 27,586 | 97.2 |
| Total | | 28,372 | - |

**Supplementary Table 12. General statistics of non-coding RNA of the genome. Related to Figure 1.**

| Type | | Number | Average length (bp) | Total length (bp) | % of genome |
|---|---|---|---|---|---|
| miRNA | | 471 | 114.66 | 54,007 | 0.01565 |
| tRNA | | 701 | 75.22 | 52,730 | 0.01528 |
| rRNA | rRNA | 133 | 196.82 | 26,177 | 0.00759 |
| | 18S | 16 | 848.25 | 13,572 | 0.00393 |
| | 28S | 26 | 127.92 | 3,326 | 0.00096 |
| | 5.8S | 9 | 138.11 | 1,243 | 0.00036 |
| | 5S | 82 | 98.00 | 8,036 | 0.00233 |
| snRNA | snRNA | 411 | 111.01 | 45,626 | 0.01322 |
| | CD-box | 277 | 97.29 | 26,949 | 0.00781 |
| | HACA-box | 48 | 130.71 | 6,274 | 0.00182 |
| | splicing | 86 | 144.22 | 12,403 | 0.00360 |
